# Efficient k-mer based curation of raw sequence data: application in *Drosophila suzukii*

**DOI:** 10.1101/2023.04.18.537389

**Authors:** Mathieu Gautier

## Abstract

Several studies have highlighted the presence of contaminated entries in public sequence repositories, calling for special attention to the associated metadata. Here, we propose and evaluate a fast and efficient *k–mer*-based approach to assess the degree of mislabeling or contamination. We applied it to high-throughput whole-genome raw sequence data for 236 Ind-Seq and 22 Pool-Seq samples of the invasive species *Drosophila suzukii*. We first used CLARK software to build a dictionary of species-discriminating *k–mers* from the curated assemblies of 29 target drosophilid species (including *D. melanogaster, D. simulans, D. subpulchrella*, or *D. biarmipes*) and 12 common drosophila pathogens and commensals (including Wolbachia). Counting the number of *k–mers* composing each query sample sequence that matched a discriminating *k–mer* from the dictionary provided a simple criterion for assignment to target species and evaluation of the entire sample. Analyses of a wide range of samples, representative of both target and other drosophilid species, demonstrated very good performance of the proposed approach, both in terms of run time and accuracy of sequence assignment. Of the 236 *D. suzukii* individuals, five were re-assigned to *D. simulans* and eleven to *D. subpulchrella*. Another four showed moderate to substantial microbial contamination. Similarly, among the 22 Pool-Seq samples analyzed, two from the native range were found to be contaminated with 1 and 7 *D. subpulchrella* individuals, respectively (out of 50), and one from Europe was found to be contaminated with 5 to 6 *D. immigrans* individuals (out of 100). Overall, the present analysis allowed the definition of a large curated dataset consisting of *>* 60 population samples representative of the worldwide genetic diversity, which may be valuable for further population genetics studies on *D. suzukii*. More generally, while we advocate careful sample identification and verification prior to sequencing, the proposed framework is simple and computationally efficient enough to be included as a routine post-hoc quality check prior to any data analysis and prior to data submission to public repositories.

## Introduction

With the democratization of sequencing technologies, the availability of genomic sequence in public repositories is increasing at an unprecedented rate. This is enabling the construction of large and highly informative combined datasets for an increasing number of model and non-model species, which in turn is refining the power and resolution of population genomics inference (e.g. 14). However, this increased availability of data comes at the cost of increased heterogeneity in the resulting combined dataset. For example, data sets may combine different sequencing library preparation protocols or technologies that are rapidly evolving with variable sequence quality or coverage. Similarly, for a given species, publicly available data may refer to original studies based on different sampling strategies consisting of either sequencing individuals (aka Ind-Seq) or pools of individuals (aka Pool-Seq) representative of some populations, the latter approach being quite popular due to its cost-effectiveness (30). Nevertheless, such technical characteristics can be taken into account in downstream analyses if an appropriate statistical framework is used.

More problematically, several recent studies have highlighted the high level of contamination in public repositories, which requires special attention when relying on the associated metadata description files (8, 11, 31). For example, working with wild-caught samples of species that are difficult to distinguish from other closely related species sharing the same habitat may lead to taxonomic errors or biological contamination of the sample. Such potential problems have already been reported in population genetic studies of *Drosophila melanogaster*, where sample contamination with *D. simulans* individuals was not uncommon (14, 19). In addition to biological sources, contamination may be of experimental (e.g., sample contamination or mislabeling) and/or computational (e.g., during data processing) origin (8). It should also be noted that these contamination problems are obviously not specific to publicly available data and may be even more pronounced in newly generated data that have not yet been analyzed.

In recent years, several software packages have been developed to assess the level of contamination in genomic datasets, which has been greatly facilitated by the active field of metagenomics. As recently reviewed by Cornet and Baurain (8), the available approaches can be classified into either database-free or reference-based methods. Database-free methods roughly consist of partitioning sequences based on their DNA composition (e.g. GC content or frequencies in short DNA sequences of a few nt), but they are not well suited for the analysis of large amounts of samples as they require a case-by-case inspection of the results (8). Reference-based methods consist of aligning sequences to a set of tagged sequences representative of all or part (e.g., genes) of the genomes of candidate species. In practice, this may allow either negative and/or positive filtering (i.e., removal of contaminating sequences or identification of sequences from some species of interest) of the sequencing data (8). To accomplish this task, approaches based on the exact matching of *k–mers* (i.e., *k* nt long DNA words) constituting the query sequences to a dictionary of labeled *k–mers* (built from target species genomes) have proven highly efficient and are now very popular for sequence taxonomic classification in the metagenomic field (24, 25, 33, 34).

Taking advantage of the high quality assemblies available for several dozen drosophilid genomes (15), the aim of this study was to rely on a *k–mer*-based approach to assess the level of contamination in public sequence data for the spotted wing Drosophila *D. suzukii. D. suzukii* originates from Asia and has recently invaded the entire European and American continents to become a major invasive insect pest causing dramatic losses in fruit production (1, 6). This species has thus become of great scientific interest, particularly to population geneticists, and several recent studies have provided informative samples for characterizing the structuring of its genetic diversity at global and whole-genome scales. Here, we focused on two recently published and publicly available Pool-Seq and Ind-Seq datasets, consisting of whole-genome sequences (WGS) for i) 22 pools of individual DNA (with n=50 to n=100 individuals per pool) representative of populations sampled both in the Asian native range (n=6) and in the European (n=8) and American invaded ranges (n=8) (22); and ii) 236 individuals collected mainly in North America but also at several sites in Europe, Brazil and Asia (18). A combined analysis of these two datasets using standard descriptive approaches revealed anomalous behavior of some samples (not shown), thus motivating a systematic screening of all samples for putative contamination or (taxonomic) misidentification problems. Indeed, as highlighted by Piper *et al*. (28), rapid morphological identification of *D. suzukii* on wild-caught specimens can be tricky. For example, the distinctive spots observed on the wing extremities are present only in (non-juvenile) males, and this feature is shared with two of its sister species *D. biarmipes* and *D. subpulchrella*, whose distributions overlap all or part of that of *D. suzukii* in its native Asian range (23, 32). In addition, both *D. suzukii* and *D. subpulchrella* females possess a large and serrated ovipositor that allows them to penetrate under the skin of ripening fruits and lay eggs (2), making the distinction between these two species even more difficult.

To assess contamination in publicly available *D. suzukii* raw sequencing data, we developed and evaluated a fast and efficient approach based on *k–mer*-based methods implemented in the software CLARK (25). We first build dictionaries of species-discriminating *k–mers* from the curated assemblies of 29 target drosophila species and 12 common drosophila pathogens and commensals. WGS data for individual samples representative of both the target and other drosophilid species were then analyzed to evaluate the performance of the proposed approaches, both in terms of run time and accuracy of sequence assignment. Finally, we analyzed publicly available WGS data for the aforementioned 236 Ind-Seq (18) and 32 Pool-Seq (22) samples of the invasive species *Drosophila suzukii*, allowing us to identify unambiguously contaminated samples.

## Material and Methods

### Construction of the Clarkand Clark-ltarget dictio-naries of species-discriminating *k–mers*

Of the 136 reference genome assemblies available for species belonging to the genus Drosophila in the NCBI repository (https://www.ncbi.nlm.nih.gov/datasets/genomes/ accessed in February 2022), 29 were retained based on assembly quality criteria such as contiguity (evaluated with contig N50) and completeness (using BUSCO scores, 20); but also and mostly based on phylogenetic criteria (Figure 1). Our goal was to obtain a good representation of species closely related to *D. suzukii*, focusing on those belonging to the two subgenera *Sophophora* and *Drosophila* that are not unambiguously resolved (see Discussion). For subgroups or groups represented by multiple species (among those with good quality assemblies available), only one target species was selected, favoring the most cosmopolitan or temperate species (13), except for the species most closely related to (and likely to be confounded with) *D. suzukii* (e.g., *D. subpulchrella* and *D. biarmipes*). To further improve the representation of *D. suzukii* in the *k–mer* dictionary, the draft assembly of Ometto *et al*. (23) was also downloaded from the ENA repository (https://www.ebi.ac.uk/ena/browser/home). Although this assembly was of lower quality than the reference (27), it was obtained from a different isofemale line and was based on short read sequences from a pool of females and males. Similarly, for *D. subpulchrella* (the sister species of *D. suzukii*), the assembly from (15) was considered in addition to the latest NCBI reference assembly (9), because it is based on male individuals and therefore contain Y-linked contigs. The high quality *D. simulans de novo* assembly from Chang *et al*. (3) was also included for similar reasons.

**Fig. 1.**
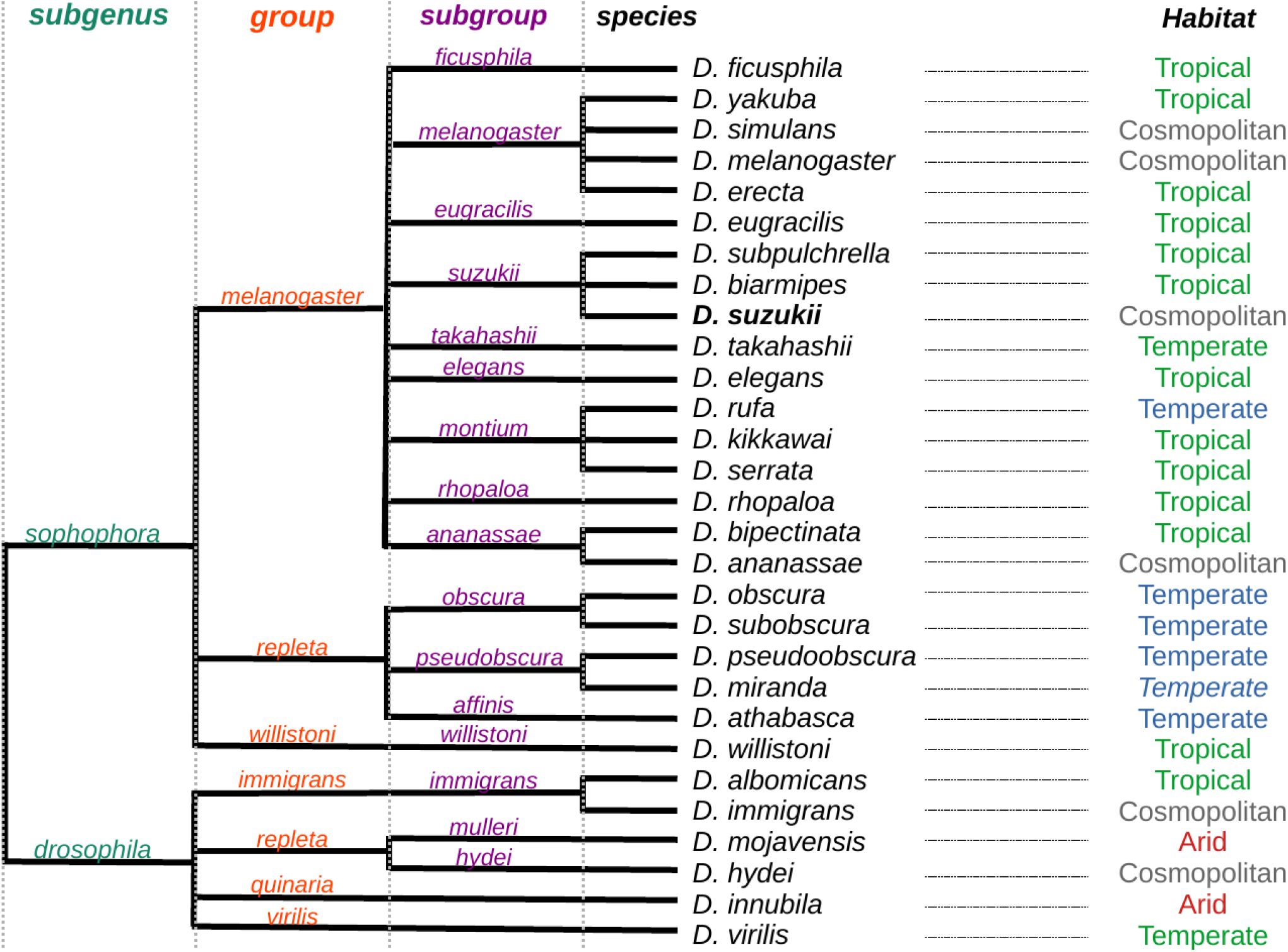
Relationship between the 29 target drosophilid species (adapted from https://www.ncbi.nlm.nih.gov/taxonomy). Species habitat was defined according to Jezovit *et al*. (13), except for *D. miranda. D. suzukii* is highlighted in bold.

The resulting 32 assemblies, described in Table 1, were further screened for non-Drosophila contaminating sequences using the program KRAKEN2 V2.1.2 (34) by querying a database constructed from the NCBI non-redundant nucleotides (nt) released in February 2020. A contig or scaffold sequence was considered contaminating if it was assigned to a taxonomic identifier unrelated to any drosophilid species. Note that contigs assigned to Wolbachia endosymbionts were also flagged as contaminating, as we chose to consider Wolbachia specifically here (see below). Of the 90,071 sequences (i.e., contigs or scaffolds) from all 32 assemblies (5.96 Gb in total), 16,123 sequences (17.9%) were found to be contaminating. As detailed in Table S1, these contaminating sequences were mostly short, ranging from 110 bp to 1,478,327 bp (median size of 1,522 bp), totaling only 102.7 Mb (i.e. 1.72% of all sequences). It should be noted that Wolbachia-related sequences represented only 6,173,139 bp of the contaminating sequences (6.01%), with the major contributor being the *D. ananassae* assembly (6,078,940 bp), which may be explained by the widespread lateral gene transfer from Wolbachia described in this species (16). The other Wolbachia contaminating sequences belonged to the assemblies for *D. suzukii* (83,189 bp) of Ometto *et al*. (23) and *D. willistoni* (11,010 bp). Finally, out of the 32 assemblies, only three (for *D. albomicans, D. innubila* and *D. melanogaster* species) were found to be free of any contaminant, the most contaminated assemblies being those for *D. immigrans* and *D. willistoni* with 10.6% and 8.1% of their total length contaminated, respectively (Table 1). As expected, the completeness of the assemblies (except for the draft assembly for *D. suzukii* from (23) mentioned above) remained quite good after filtering out contaminating sequences, with more than 98% (resp. 95%) of the 3,285 BUSCO genes of the `diptera_odb10` dataset (20) identified in 26 (31) of the assemblies (Table 1). Finally, in addition to the drosophilid species and following Kapun *et al*. (14), 13 genome assemblies representing twelve different common drosophilid commensals and pathogens were included in the construction of the *k–mer* dictionaries (Table 1). Note that the two reference assemblies for the Wolbachia endosymbiont of D. melanogaster and D. simulans were used to represent Wolbachia.

**Table 1.**
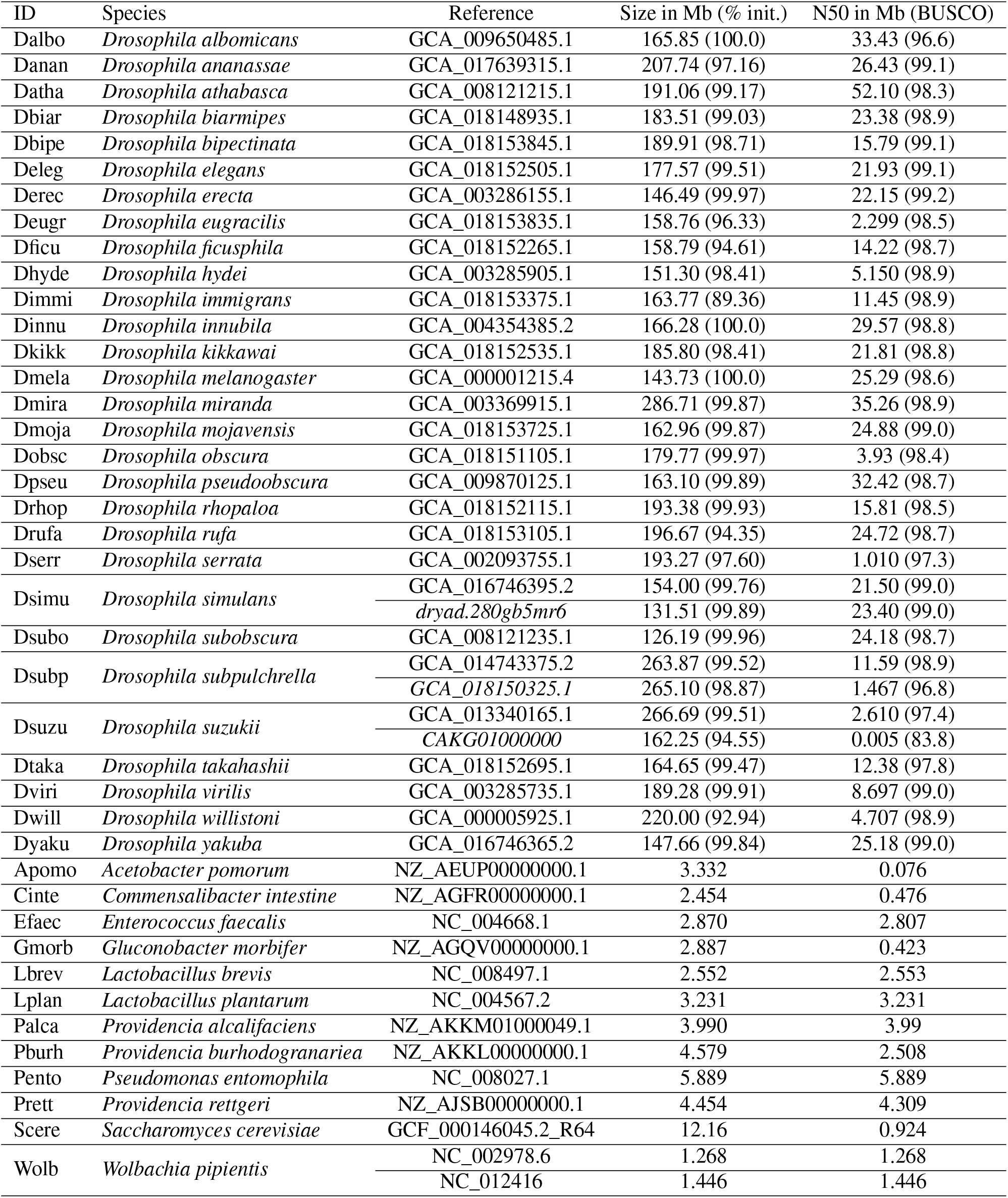
Description of the reference genome assemblies for the 29 drosophilid species (n=32 assemblies) and 12 common commensals and pathogens (n=13 assemblies) used to build the target *k–mer* dictionaries. All genome assemblies were downloaded from the NCBI repository (https://www.ncbi.nlm.nih.gov), except for the two additional assemblies for *D. simulans* and *D. suzukii*, which were downloaded from the Dryad (https://datadryad.org) and ENA (https://www.ebi.ac.uk/ena) repositories, respectively (with accession ID in italics in the third column). The size and N50 of all assemblies are given in the fourth and fifth columns. For drosophilid species, these correspond to the assemblies after filtering out the identified contaminant contigs (or scaffolds), the percentage of the original assembly retained is given in parentheses in the fourth column. Similarly, the BUSCO scores in parentheses correspond to the percentage of complete genes identified among the 3,285 genes of the `diptera_odb10` dataset (20).

From the 45 reference assemblies representing the 29 drosophilid, commensal, and pathogen species, two different dictionaries of species-discriminating *k–mers* (i.e., *k– mers* that occur exclusively in the genome of a species represented by one or more assemblies) were then constructed using versions 1.2.6.1 of CLARK (default k=31) and Clark-L (default k=27), respectively (25). Clark-lis a variant of CLARK designed for use when the amount of RAM is limited, with minimal impact on assignment accuracy. Both CLARK and Clark-lwere run in single-threaded mode on a computer cluster grid. Building the *k–mer* dictionary (on a single thread of a cluster node equipped with a processor Intel^®^ Xeon^®^ CPU E5-2683 v4 @2.10GHz) took 2h46min with a peak RAM usage of 128G using CLARK and 55s with a peak RAM usage of 2.65G using Clark-l. The resulting database consisted of 3,714,249,662 31-mers and 50,311,519 27-mers, respectively, and required 47.8 Gb and 1.97 Gb of RAM to load when computing the query sequence classification with CLARK and Clark-l, respectively.

### Query short-read sequencing data

A total of 301 short read WGS data sets were downloaded from the public SRA repository (https://www.ncbi.nlm.nih.gov/sra). These include 43 samples used for the empirical evaluation of *k–mer*-based assignment accuracy, derived from the sequencing of laboratory strains representative of different drosophilid species (including data on 12 of the 29 target species available for the strains used to generate the corresponding assemblies) and the Wolbachia endosymbiont of *D. melanogaster* (Table S2). As detailed in Table S2, all of these data were obtained from paired-end (PE) sequencing (2 *×* 150 nt) on an Illumina HiSeq 4000 instrument, except for ten samples sequenced on an Illumina i) GAIIX in PE125 mode (n=1); ii) NextSeq550 in PE150 mode (n=4); iii) HiSeq 2000 in PE100 (n=2) and PE150 (n=1) modes; iv) MiSeq in PE300 mode (n=1); or v) HiSeq Ten X in PE150 (n=1). The second type of data corresponded to WGS data for 236 *D. suzukii* individuals (Ind-Seq data) representative of 40 population samples (4-10 ind. per sample, mean=5.9) published by Lewald *et al*. (18). These samples were mainly collected in the continental USA (n=31). The other regions represented are Brazil (n=1); Europe (n=2; Ireland and Italy) for two of them; China (n=2); South Korea (n=2), but also Japan (n=1) and Hawaii (n=1), via two laboratory strains. These were all sequenced on an Illumina HiSeq4000 in PE150 (n=201) or PE100 (n=35) mode (Table S3). The last type of data corresponded to WGS data from 22 pools of *D. suzukii* individuals (Pool-Seq data) representing 22 worldwide populations representative of the Asian native range (n=6) and the European (n=8) and American (n=8) invaded ranges, published by Olazcuaga *et al*. (22). These were all sequenced on an Illumina HiSeq2500 in PE125 mode (Table S3).

Raw PE reads were filtered with `fastp` 0.23.1 (4) with the default options to remove contaminating adaptor sequences and trimmed for poor quality bases (i.e. with a phred quality score <15). In addition, the `--merge` and `--include_unmerged` options were used to merge the detected overlapping PE reads into a single sequence. Finally, the `--stdout` option was enabled to generate an interleaved `fastq` output, which was converted to `fasta` format (losing quality and pairing information) with a simple `awk` one-liner for assignment analysis. As shown in Figure S1 and detailed in Tables S2 and S3, the quantity (and quality) of sequencing data was highly variable between samples, with the percentage of non-overlapping sequences ranging from 5.79 to 94.4 (median 35.0) as a consequence of different insert sizes; and the estimated percentage of duplicate reads ranging from 0.69 to 24.8 (median 4.44) (Figure S1B). Note that the sequencing data were not de-duplicated here, although this may be possible using the latest version of `fastp` (4).

### Assignment of query sequences and contamination estimation

For each sample, the sequences contained in the filtered `fasta` files were matched to the target dictionaries of species-discriminating *k–mers* using Clark and Clark-L (25). Briefly, analyzing a sequence consists of first decomposing it into its constituent *k–mers* (i.e., a *L* nt long sequence can be decomposed into *L*− *k* + 1 *k–mers* of length *k* nt) of length *k* = 31 and *k* = 27 for CLARK and Clark-l, respectively. Each *k–mer* is then searched in the corresponding target dictionary and, if found, assigned to the underlying target species. Counting the number of *k–mers* assigned to the different species then provides a simple decision criterion for sequence classification. More specifically, for a given sequence, let *t*_1_ and *t*_2_ be the target species with the highest and second highest counts (*k*_*q*_(*t*_1_) and *k*_*q*_(*t*_2_) *≤ k*_*q*_(*t*_1_)) of matching *k–mers*. If no species-discriminating *k–mer* was found in the sequence (i.e., *k*_*q*_(*t*) = 0 for all target species *t*), the sequence is unassigned. If *k*_*q*_(*t*_1_) *>* 0, the sequence is assigned to species *t*_1_ with a ‘confidence score’ defined as 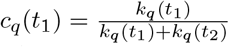, noting that *c*_*q*_(*t*_1_) = 1 if all the matching *k–mers* are assigned to *t*_1_ (i.e., *k*_*q*_(*t*) = 0 for all *t* ≠ *t*_1_). At the sample level, the origin and level of contamination can then be further assessed by counting the number of sequences assigned to the different target species. In practice, CLARK was run with option `-s 2` to load only half of the species-discriminating *k–mers* in the target dictionary, following the manual recommendation indicating that this value ‘represents a good trade-off between speed, accuracy and RAM usage’. Both CLARK and Clark-lwere run with the options `-n 1` (i.e., on a single thread) and `-m 0` (to compute the confidence score). The resulting `csv` files were parsed with a custom `awk` script to count for each sample i) the total number of sequences with no matching *k–mer*; ii) the total number of sequences with at least one matching *k–mer*; and iii) the proportion of sequences assigned to each target species. Four different criteria were considered for assigning sequences to their inferred species *t*, taking into account both the minimum number *nk*_min_(*t*) of matching *k–mers* and the confidence score *c*_*q*_(*t*): i) *nk*_min_(*t*) *≥* 1 and *c*_*q*_(*t*) *>* 0.9; ii) *nk*_min_(*t*) *≥* 1 and *c*_*q*_(*t*) *>* 0.95; iii) *nk*_min_(*t*) *≥* 5 and *c*_*q*_(*t*) *>* 0.9; and the most stringent iv) *nk*_min_(*t*) *≥* 5 and *c*_*q*_(*t*) *>* 0.95. All subsequent analyses were performed using the `R` software (29).

## Results

### Clark and Clark -l run times

Publicly available short-read WGS data for 301 different samples derived from i) laboratory strains representing different drosophilid species (n=43); ii) 236 (putative) *D. suzukii* individuals representing 40 different populations; and iii) 22 pools of *D. suzukii* individuals representing 22 different populations were assigned to two different species-discriminating *k–mer* dictionaries built from the curated assemblies available for 29 drosophilid species (Figure 1) and 12 common drosophila commensals and pathogens (Table 1), using the *k–mer*-based approaches implemented in CLARK and Clark-l(25). Although this step is not required for assignment, the raw PE reads were filtered to limit the potential impact of varying sequence quality on the assessment of assignment efficiency and accuracy, particularly with respect to the observed proportion of unassigned sequences per query sample. After filtering, the total number of sequences per sample ranged from 1.61 × 106 to 367 × 106 (median of 18.5*×* 106) for a total length ranging from 0.248 Gb (i.e. ∼ 0.9X of the *D. suzukii* genome) to 36.9 Gb (i.e. ∼ 137X of the *D. suzukii* genome). The sequence length was representative of typical short read datasets, with a sample mean length (after merging overlapping reads) ranging from 92.7 bp to 287 bp (Figure S1C).

Tables S2 and S3 show the total Clark and Clark-lrun times *t*_*r*_ for each sample, together with the time *t*_*l*_ required to load the corresponding *k–mer* target dictionary and the time *t*_*a*_ required to assign all sequences (*t*_*r*_ = *t*_*l*_ + *t*_*a*_). As summarized in the Table 2, *t*_*l*_ was a few seconds for Clark-land a few minutes for C<sc>LARK</sc>, the C<sc>lark-L</sc> target dictionary containing about 75 times less *k–mer* than C<sc>LARK’s</sc> (see M&M). In addition, Clark-lrequired much less RAM than CLARK (1.97 Gb *vs* 47.8 Gb), allowing it to run on a standard laptop. Note that CLARK and Clark-lwere run sequentially on each sample on a computer grid, but the samples were analyzed in parallel. Therefore, the run times between samples may be somewhat dependent on the characteristics of the underlying node, which explains the observed variation in dictionary loading times.

**Table 2.**
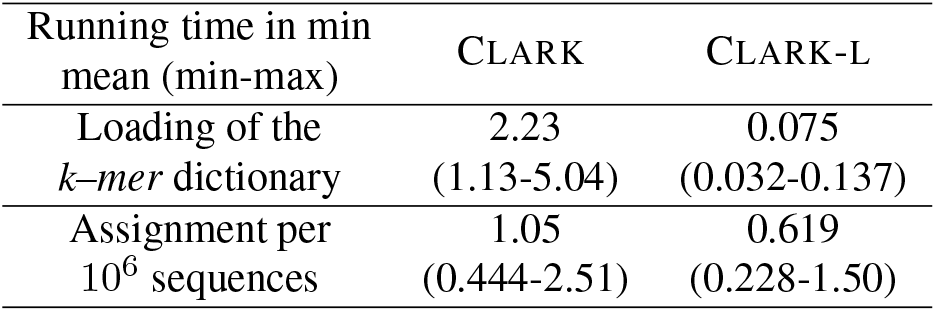
Mean Clark and Clark-lrun times (minimum-maximum) across the analyses of the 305 short-read sequencing datasets. Each analysis was run on a single thread of a cluster node equipped with a processor Intel^®^ Xeon^®^ CPU E5-2683 v4 @2.10GHz

| Running time in min<br>mean (min-max) | CLARK | CLARK-L |
| --- | --- | --- |
| Loading of the<br>*k*-mer dictionary | 2.23<br>(1.13-5.04) | 0.075<br>(0.032-0.137) |
| Assignment per<br>10<sup>6</sup> sequences | 1.05<br>(0.444-2.51) | 0.619<br>(0.228-1.50) |

Given the size of the data sets, most of the analysis time was spent on sequence assignment which was almost linearly related to the number of sequences (Figure S2) as sequence length was similar across samples (Figure S1C). On average, the analysis of 1 million sequences (i.e., ∼ 0.56X of the *D. suzukii* genome with 150 nt reads) took 0.619 and 1.05 minutes with Clark-land CLARK, respectively (Table 1), making both approaches highly computationally efficient.

### Proportion of assigned sequences

The percentage of sequences with no matching *k–mer* (i.e., not assignable) was similar between CLARK (ranging from 2.29% to 85.5% with a median value of 20.1%) and Clark-l(ranging from 4.07% to 86.1% with a median value of 15.7%) (Figures 2A and 2B). Surprisingly, this percentage tended to be slightly lower for the *D. suzukii* sample (Ind-Seq or Pool-Seq) when analyzed with Clark-l, which may be related to the smaller *k–mer* size (k=27 for Clark-land k=31 for Clark) leading to lower specificity. However, the proportion of sequences with no matching *k–mer* remained higher for Clark-lanalyses for samples representative of the other species either represented or not represented in the target dictionaries (Figure 2B). As expected, and regardless of the program used, the highest percentages were observed for samples belonging to species not represented in the target dictionaries (up to 85.5% and 86.1% of sequences with no matching *k–mer* for the *D. repleta* sample analyzed with Clark and Clark-l, respectively), although the distribution was very wide and almost bimodal due to some samples being represented by closely related target species (see below). The sample representing target species had the lowest number of sequences with no matching *k–mer*, most of them (including *D. suzukii*) corresponding to short-read sequence data obtained from the same strains used to generate the reference assembly, with the notable exception of *D. melanogaster, D. simulans*, and the Wolbachia sample (see below), which were also outliers in the distributions of Figure 2B (see Table S4). Their values was actually similar to wild-caught *D. suzukii* samples (see below). The *D. simulans* sample was obtained from Madagascar individuals (26) thus distantly related to the two reference assembly strains, which may explain the observed pattern (see Discussion). Likewise, the analyzed *D. melanogaster* sample corresponded to a pool of 162 isogenic strains from the DGRP panel and may thus display higher genetic diversity (35).

**Fig. 2.**
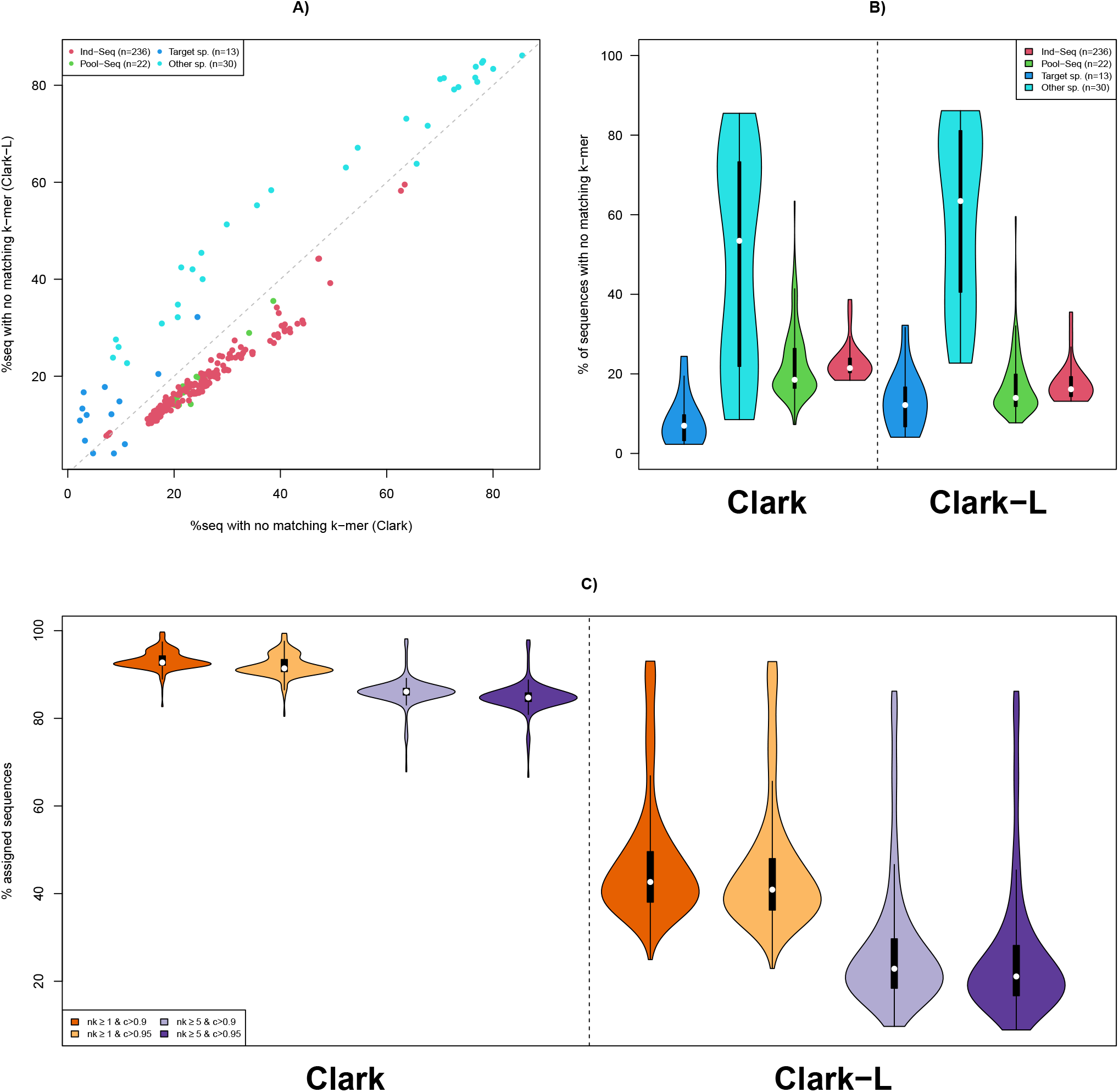
Sequence assignment rate for the 301 samples analyzed with Clark and Clark-l. A) Percentage of sequences with no matching *k–mer* in the target dictionaries. Samples are colored according to their origin, i.e. i) dark blue if from species represented in the target dictionary (‘Target sp.’); ii) light blue if from drosophilid species not represented in the target dictionary (‘Other sp.’); iii) green for *D. suzukii* individuals from Lewald *et al*. (18) (‘Ind-Seq’); and iv) red for pools of *D. suzukii* individuals from Olazcuaga *et al*. (22) (‘Pool-Seq’). B) Violin plots showing the distribution of the percentage of sequences with no matching *k–mer* in the corresponding target dictionary with Clark (left panel) and Clark-l(right panel) analyses. For each analysis, four distributions are shown for the different sample origins (same color code as in A). C) Distribution of the percentage of assigned sequences (among those with at least one species-discriminating *k–mer* from the target dictionary) for four filtering criteria on i) the number *nk* of matching *k–mers* (*nk ≥* 1 or *nk ≥* 5); and ii) the assignment confidence score *c* as defined in the main text (*c >* 0.9 or *c >* 0.95).

Consistent with a lower specificity of Clark-l(suggested by the unexpectedly slightly lower proportion of sequences with no matching *k–mer* in *D. suzukii* individuals), the percentages of assigned sequences among assignable sequences (i.e., containing at least one *k–mer* matching the dictionary of target species discriminating *k–mers*) were much lower with Clark-lthan with CLARK (Figure 2C). The percentages of assigned sequences always decrease with the stringency of the filtering criteria on the number *nk* of matching *k–mers* (*nk* ≥ 1 or *nk* ≥ 5) and the assignment confidence score *c* (as defined above in the M&M section), with the threshold on *nk* having the strongest effect. At the most stringent criterion (*nk ≥* 5 and *c* > 0.95), which was chosen for the remainder of this study, 84.8% and 26.4% of sequences with at least one matching *k–mer* were assigned to CLARK and Clark-l, respectively, on average (see Tables S4 and S5 for details).

### Assignment accuracy for samples representative of target and other species

To empirically evaluate the extent to which the proportion of assigned sequences from a sample provides an accurate proxy for species assignment, we focused on the results obtained for the 13 short-read datasets derived from strains representative of one of the target species (including Wolbachia), but also on 30 additional samples representative of unrepresented drosophilid species, considering our most stringent filtering threshold for sequence assignment (Figures 3 and S3 for CLARK and Clark-l results, respectively). The results obtained were highly consistent for all 13 samples representing the target reference species. More precisely, with CLARK, the percentage of sequences assigned to their species of origin was >99% or close to 99% (with 98.9% for both *D. suzukii* and *D. subpulchrella*) for 9 of these samples. The remaining four were those belonging to i) *D. biarmipes* (94.0%), due to yeast contamination with 5.76% of the sequences assigned to *S. cerevisiae*; ii) *D. melanogaster* (94.1%) with 3.78% of the analyzed sequences assigned to Wolbachia and 1.11% to *D. simulans*; iii) *D. simulans* (93.6%) with 3.14% of the analyzed sequences assigned to Wolbachia and 2.54% to *D. melanogaster*; and iv) Wolbachia with only 5.58% actually assigned to Wolbachia and 94.0% to *D. melanogaster* (Table S3). Note that this latter Wolbachia sample was actually obtained from sequencing a *D. melanogaster* strain, and the observed level of contamination was in close agreement with the 5% of reads mapping to the Wolbachia wMel genome by the original authors (21). Similar results were obtained when scanning these 13 samples with Clark-l (Figure S3 and Table S5), with some notable differences. Indeed, the percentage of sequences assigned to their species of origin was also above 99% (including the *D. subpulchrella* one) or close to it (with 98.0% for *D. yakuba*) for 8 of the 9 samples that showed similarly high assignment rates with CLARK. However, it was substantially lower for the *D. suzukii* sample (92.1%), with 7.22% of its sequences assigned to the *D. subpulchrella* sister species. Similarly, only 86.2% of the *D. melanogaster* sample sequences were assigned to *D. melanogaster*, with 6.86%, 2.62%, 1.65%, and 1.40% assigned to *D. suzukii, D. simulans, D. virilis*, and Wolbachia, respectively. Conversely, the percentage of correctly assigned sequences was higher with Clark-l than with CLARK for the *D. biarmipes* (96.0%); *D. simulans* (98.1%) and Wolbachia (40.0% with 55.8% assigned to *D. melanogaster*) samples, the latter apparently being overestimated.

**Fig. 3.**
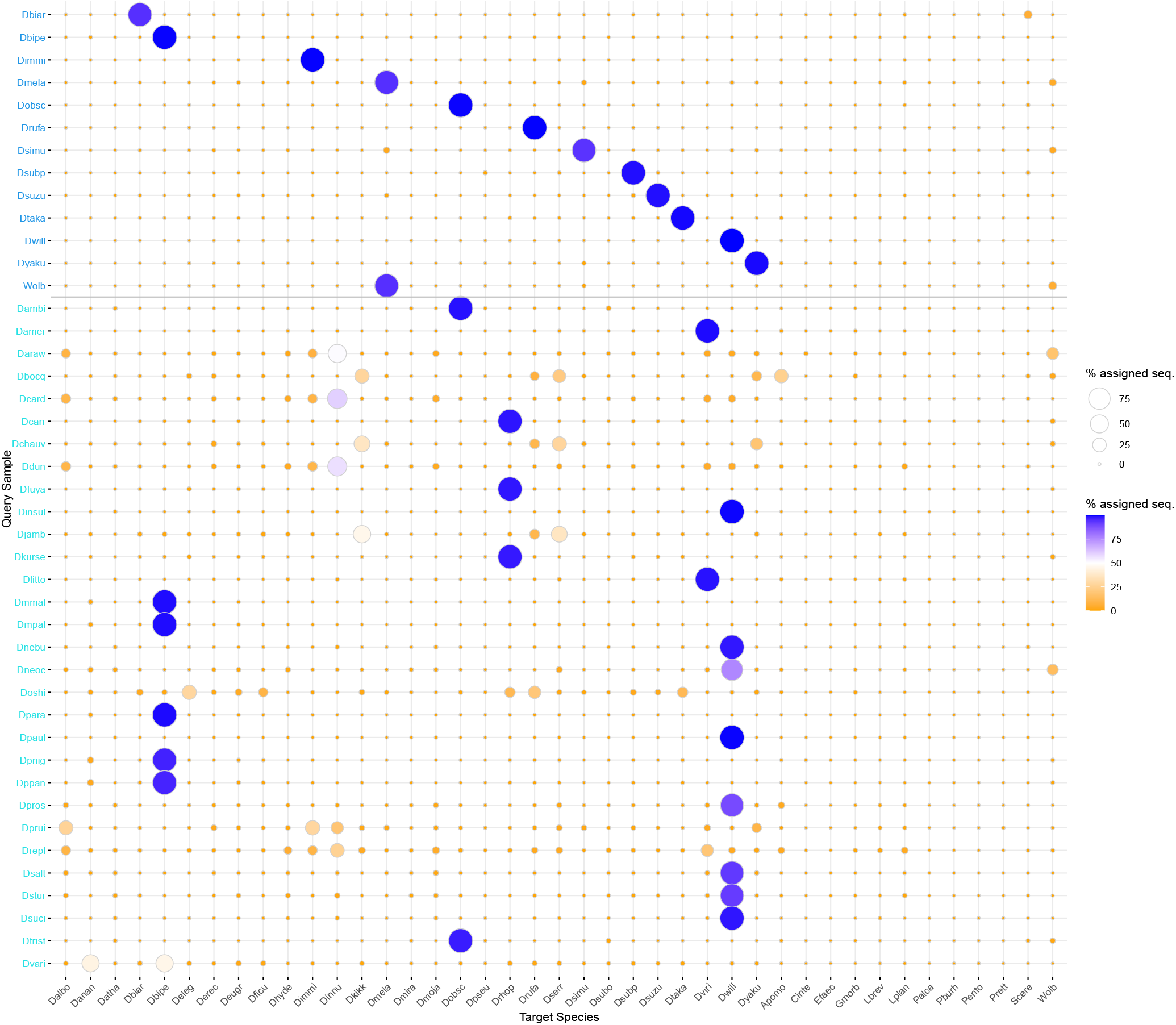
Bubble plots summarizing assignment results obtained with Clark using the most stringent sequence assignment criterion (i.e., *nk ≥* 5 and *c >* 0.95, see the main text) for 13 samples (labeled in dark blue on the top of the y-axis) belonging to target species represented in the target *k–mer* dictionary and 30 other unrepresented drosophilid species.

Of the 30 samples from non-target species, 16 had more than 96% of their reads assigned to a single target species by CLARK (Figure 3). As expected, the corresponding species was generally the most closely related (15). More precisely, samples from i) *D. paulistorum* and *D. insularis* (*D. willistoni* subgroup) and *D. sucinea* and *D. nebulosa* (*bocainensis* sub-group from the *willistoni* group) had 99.7%, 99.7%, 98.1%, and 97.9% of their sequences assigned to *D. willistoni*, respectively; ii) *D. parabipectinata, D. malerkotliana pallens, D. malerkotliana malerkotliana, D. pseudoananassae*, and *D. pseudoananassae nigrens*, all of which belong to the *ananassae* subgroup, had 99.2%, 99.1%, 99.0%, 96.5%, and 96.0% of their sequences assigned to *D. bipectinata* (*ananassae* sub-group), respectively; iii) *D. ambigua* and *D. tristis* (*obscura* subgroup) had 98.7% and 97.3% of their sequences assigned to *D. obscura*, respectively; iv) *D. americana* and *D. littoralis* (*virilis* group) had 99.2% and 98.6% of their sequences assigned to *D. virilis*, respectively; and finally v) *D. carrolli, D. fuyamai*, and *D. kurseongensis* (*rhopaloa* subgroup) had 98.2%, 98.0%, and 97.7% of their sequences assigned to *D. rhopaloa*, respectively. As shown in Figure S4A, these 16 samples also had percentages of sequences with no matching *k–mer* in the range of those observed for samples from target species (Figure 2), i.e. *<*40% except for *D. sucinea* and *D. nebulosa*. For the other samples from the most distantly related species, both the highest observed assignment rate (to a target species) and the percentage of sequences with no matching *k–mer* clearly suggested that the target repository was not representative. At the extreme, the most represented target species capture less than 30% of the assigned sequences for the samples from *D. repleta, D. pruinosa, D. ohnishii*, and *D. bocqueti* (Figures 3 and S4A). Such species may therefore be considered unassignable with the current version of the *k–mer* dictionary. Despite a higher proportion of sequences with no matching species-discriminating *k– mer*, very similar results were obtained with Clark-l (Figure S3 and S4B).

### Scanning 236 Ind-Seq and 22 Pool-Seq *D. suzukii* WGS data

As summarized in Figure 4 (see Table S4 for details), sequences from the 236 Ind-Seq (18) and 22 Pool-Seq (22) *D. suzukii* were generally assigned to *D. suzukii* by CLARK. More precisely, 215 of the 236 Ind-Seq and 17 of the 22 Pool-Seq showed > 95% of their (assigned) sequences assigned to *D. suzukii*, with a median proportion of 97.5% over the 258 samples. It should be noted that these 215 individuals and 17 pools, which can be unambiguously considered as fully *D. suzukii*, all had a non-negligible fraction of their sequences assigned to *D. subpulchrella* with a median of 1.94% (ranging from 1.50% to 3.12%) and 2.20% (ranging from 1.96% to 2.64%), respectively. These proportions were higher than the one observed for the *D. suzukii* reference sample (0.433%) and may be related to the incomplete representation of genetic diversity within *D. suzukii* by the *k–mer* dictionary (see Discussion). Conversely, the results allowed 16 clearly mislabeled *D. suzukii* individuals to be identified as *D. simulans* (n=5) or *D. subpulchrella* (n=11). These consist of i) the 5 individuals (with US-Ca2 ID prefix, Table S2) sampled simultaneously in Watsonville (California, USA) with 92.1% to 96.9% of their sequences assigned to *D. simulans* (96.9% to 98.4% if Wolbachia is also included); ii) the 5 individuals (with Ko-Nam ID prefix, Table S2) sampled in Namwon (South Korea) with 97.9% to 98.7% of their sequences assigned to *D. subpulchrella*; iii) one of the 10 individuals (with Ko-San ID prefix, Table S2) sampled in Sancheong (South Korea) with 96.4% of its sequences assigned to *D. subpulchrella* (the other 9 individuals showing only 1.71% to 2.09% of their sequences assigned to *D. subpulchrella*); and iv) four of the five individuals (with CN-Kun ID prefix, Table S2) sampled in Kunming (Yunnan, China) with 97.3% to 97.6% of their sequences assigned to *D. subpulchrella*. The last CN-Kun individual had a unique pattern with 88.1% of its sequences assigned to *D. subpulchrella* and 9.58% assigned to *D. suzukii*, which may be consistent with a recent hybrid origin (see Discussion). For the 10 individuals that can be unambiguously considered as fully *D. subpulchrella* (i.e. with >95% of their sequences assigned to *D. subpulchrella*), an assignment pattern opposite to that of the *D. suzukii* individuals was observed, as all of them had a non-negligible fraction of their sequences assigned to *D. suzukii* with a median value of 1.61% (ranging from 1.14% to 2.78%).

**Fig. 4.**
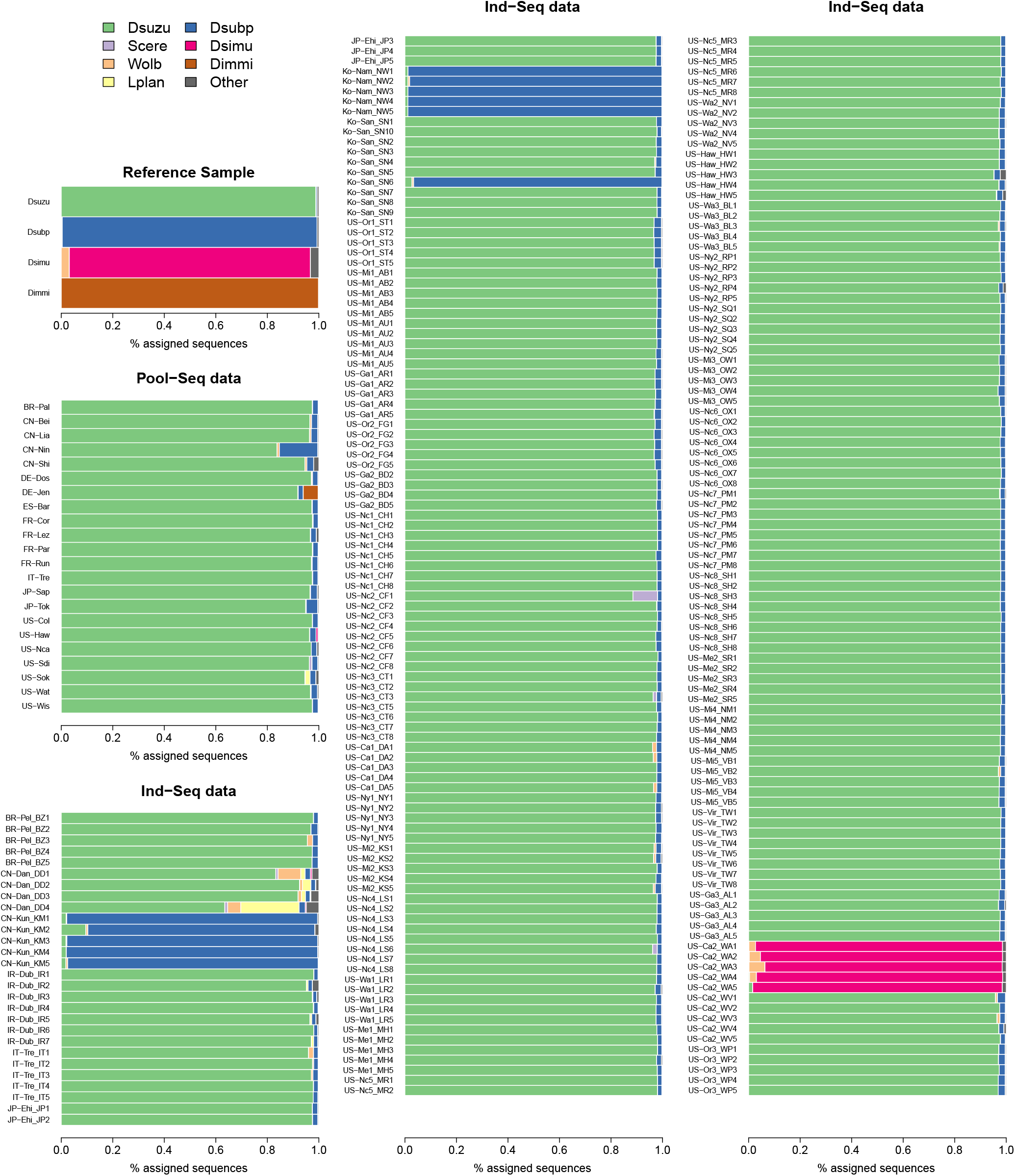
Barplots summarizing assignment results obtained with Clark using the most stringent sequence assignment criterion (i.e., *nk ≥* 5 and *c >* 0.95, see the main text) for the *D. suzukii* Ind-Seq (n=236) and Pool-Seq (n=22) samples. For each sample, the proportions of sequences assigned to the 7 target species that contribute at least 5% of the sequences of one of any of the 258 samples are shown using the color code indicated in the top-left legend. The proportions of sequences assigned to the 34 other target species are shown in gray.

Among the 22 Pool-Seq samples, two to three pools were found to be likely contaminated with non-*D. suzukii* individuals. These are i) the DE-Jen pool of 100 individuals sampled in Jena (Germany), which contains 5.79% of sequences assigned to *D. immigrans*; ii) the CN-Nin pool of 50 individuals sampled in Ningbo (Zhejiang, China), which contains 15.0% of sequences assigned to *D. subpulchrella* (and 83.8% to *D. suzukii*); and iii) the JP-Tok pool of 50 individuals sampled in Tokyo (Japan), with 4.47% of sequences assigned to *D. subpulchrella* (and 94.9% to *D. suzukii*). Assuming an equal contribution of pool individuals to the Pool-Seq sequences, the DE-Jen pool may actually contain up to 6 *D. immigrans* individuals (and 94 *D. suzukii* individuals). Furthermore, to estimate the number of *D. subpulchrella* individuals in contaminated pools while accounting for *D. suzukii* and *D. subpulchrella* cross-assignment of sequences, let 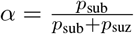 be the relative proportion of sequences assigned to *D. subpulchrella*. Based on the median proportions observed in the Ind-Seq samples, the following rough estimates were obtained: 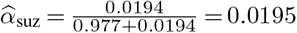 for *D. suzukii* individuals and 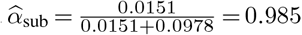 for *D. subpulchrella* individuals. The number of *D. subpulchrella* individuals *n*_sub_ in a contaminated pool of *n* individuals can then simply be derived from these estimates using their observed relative proportion *α*_*o*_ as 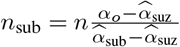. This leads to an estimated number of *D. subpulchrella* individuals of 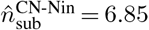 and 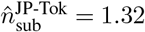 i.e. probably 7 and 1 *D. subpulchrella* individuals within the CN-Nin and JP-Tok pools, respectively. Overall, very low levels of Wolbachia contamination were detected within the Ind-Seq and Pool-Seq samples, with median proportions of assigned sequences of 3.80 × 10*−*4% and 0.145%, respectively. However, 14 samples (Ind-Seq only) had more than 1% of their sequences assigned to Wolbachia. They consisted of i) the five US-Ca2 individuals mentioned above, which are actually *D. simulans*, with proportions ranging from 1.08% to 6.17%; ii) the four individuals with the CN-Dan ID prefix (Table S2), sampled in Dandeong (China), with proportions ranging from 1.07% to 8.82%; iii) three of the five individuals with the US-Ca1 ID prefix (Table S2) sampled in Davis (California, USA) with proportions ranging from 1.29% to 1.55%; iv) one of the five individuals with the BR-Pel ID prefix sampled in Pelotas (Brazil) with a proportion of 2.07%; and v) one of the five individuals with the IT-Tre ID prefix sampled in Trento (Italy) with a proportion of 1.92%. Finally, a few Ind-Seq and Pool-Seq samples showed non-negligible to substantial contamination with five of the 11 other microbial species represented in the *k–mer* target dictionary. For example, more than 1% of the sequences were assigned to the *L. plantarum* bacterial gut symbiont for five samples corresponding to i) the four CN-Dan individuals (see above), with proportions ranging from 1.55% to 22.6%; and ii) the US-Sok pool of 50 individuals sampled in Dayton (Oregon, USA) with a proportion of 1.87%. Similarly, > 1% of the sequences were assigned to *S. cerevisiae* yeast for five samples corresponding to i) one of the four CN-Dan individuals with a proportion of 1.12%; ii) three individuals (with ID prefixes US-Nc2, US-Nc3, and US-Nc4, Table S2) sampled in different locations in North Carolina (USA) with proportions ranging from 1.33% to 9.58%; and iii) the US-Sdi pool of 50 individuals sampled in San-Diego (California, USA) with a proportion of 1.02%. At the margin, three other microbial species were also found to be represented by more than 1% of the sequences in at least one sample. These are i) the *A. pomorum* gut bacteria in two Chinese (CN-Dan) individuals (with proportions of 1.04% and 1.46%) and in the CN-Shi pools of 50 individuals sampled in Shiping (China) with 1.56%; ii) the *L. brevis* intestinal bacteria also found in two Chinese (CN-Dan) individuals with proportions of 1.12% and 4.19%; and iii) the *E. faecalis* pathogens in an Irish individual with proportion of 1.65%.

As expected from the assignment of *D. suzukii* and *D. subpulchrella* reference samples, Figure S5 (see Table S5 for details) suggested a worse performance of Clark-l. The proportions of *D. suzukii* sequences appeared to be substantially underestimated, with a higher effect of cross-assignment with *D. subpulchrella*. In addition, Clark-ldid not allow to detect the presence of the microbial target species as detected by CLARK.

## Discussion

The primary objective of this study was to propose and evaluate a computationally fast and accurate method for assessing contamination levels in publicly available WGS data for the *D. suzukii* species, which has been increasingly studied over the past decade. The availability of high quality genome assemblies for a wide range of drosophilid species (15) made it possible to rely on a *k–mer*-based approach consisting of constructing and querying dictionaries of species-discriminating *k–mers*. Such an approach has already proven to be quite valuable and benefits from the availability of optimized software, such as KRAKEN (33, 34) or CLARK (24, 25), which were primarily developed for metagenomics applications but have also been proposed for contaminant detection (8). As in the latter case, our primary goal here was to classify sequences at the level of predefined (target) species, and CLARK thus seemed particularly attractive due to its computationally efficient, tractable, and flexible way of both constructing and querying user-defined *k–mer* dictionaries. Although KRAKEN may be able to further assign higher-level taxonomic labels by considering phylogenetic relationships among target species, this feature was not critical for our purpose. In fact, it may have made it more difficult in practice, since the phylogeny among Drosphilidae species is far from being fully and unambiguously resolved. In particular, Finet *et al*. (10) recently provided evidence for a paraphyletic status of the subgenus Sophophora, to which most of the target species belong (Figure 1). However, as illustrated by the assignment of sequences from species closely related to one of the represented groups or subgroups (e.g., *ananassae* or *obscura*) but not included in the construction of the *k–mer* dictionary, species-level assignment provided consistent results about their origin. Yet, assignment of samples to species belonging to groups or subgroups less well represented by the target species should be interpreted with caution, especially when the observed proportion of non-matching *k–mers* is high (Figure S4). In such cases, analysis with a newly built *k–mer* dictionary including more closely related species may be valuable. Indeed, our main focus was on the evaluation of *D. suzukii* samples. We therefore chose to deliberately overrepresent the *suzukii* subgroup in the *k–mer* dictionary construction by including the high quality genome assemblies available for *D. suzukii, D. subpulchrella*, and *D. biarmipes*. The latter two species were in fact the most likely confounders in field-collected samples from the Asian range of *D. suzukii* (see Introduction). Interestingly, the inclusion of these closely related species seemed to have only a limited effect on the number of discriminating *k–mers* in the resulting dictionary, with the percentage of sequences with no assigned *k–mer* for their corresponding reference samples being in the range of that observed for reference samples from other target species (Figure 2).

Searching the resulting *k–mer* dictionary of target species sequences with CLARK (25) was highly efficient in terms of both run time and memory requirements. This makes analyses of common short-read sequencing data tractable on standard workstations or computer grids, and even on a standard laptop when using the lighter CLARK version (25), although at some moderate cost in assignment accuracy. More specifically, it took only a few minutes and about 50 Gb of RAM to load the CLARK dictionary (<1 min and <2 Gb of RAM for the Clark-ldictionary), and the mapping took about one minute per million of typical 150 nt short reads. Such assignment analyses could thus be performed routinely and may be worth including as a standard part of the quality control of sequencing data, at least for the *D. suzukii* sample. Note that here we have chosen to screen sequences after filtering raw PE reads with `fastp` (4), primarily to limit the potential impact of varying sequence quality across samples on the assessment of assignment accuracy (e.g., proportion of sequences assigned). Although this is not required in practice when trying to assign samples or assess their contamination levels, it seems to be a reasonable strategy when combined with other quality control procedures. Finally, for contamination assessment at the whole-sample level, *k–mer*-based approaches represent an attractive and efficient mapping-free alternative to competitive mapping methods that consist of mapping sequencing reads to hologenomes constructed from target species assemblies (e.g. 14). It also allows for easy interrogation of a wider range of target species, providing good quality genome assemblies are available. For sequence filtering purposes, however, such approaches must be used with caution because they rely on species-discriminating *k– mers* and thus may leave a substantial fraction of sequences unassigned. More advanced (and computationally expensive) methods may then be valuable, such as the one implemented in CLARK-S (24), which allows some mismatches in *k–mer* matching to improve the sensitivity of sequence assignment or even KRAKEN (33, 34), which was used here to identify contaminating contigs in the assemblies of the target species. Indeed, this program can rely on *k–mers* shared by several species for sequence assignment, and not only species discriminating *k–mers*, since all the *k–mers* of the target dictionary (possibly built from very large databases such as the NCBI nt) are mapped to the nodes of a phylogenetic tree (species discriminating *k–mers* to terminal nodes and shared *k–mers* to internal nodes).

Overall, the results obtained from the analysis of WGS data for reference samples belonging to different target species and single or pools of *D. suzukii* individuals demonstrated the high accuracy of the *k–mer*-based approach. It also allowed the unambiguous identification of 16 mislabeled *D. suzukii* individuals among the 236 (i.e. 6.78%) from the Lewald *et al*. (18) study. Five corresponded to *D. simulans* individuals collected at the same site in Watsonville (California, USA). It should be noted that Lewald *et al*. (18) discarded these samples from their analysis because they displayed too low mapping rates like the Dandong (China) sample, which was found here to be substantially contaminated with microbial (and Wolbachia) sequences. The eleven other non-*D. suzukii* individuals from three different locations in Asia could all be assigned to *D. subpulchrella* individuals. These were also identified as *D. subpulchrella* by Lewald *et al*. (18) (and discarded from their analysis) using a phylogenetic analysis of the mitochondrial COX2 gene. Two of the 22 Pool-Seq samples of (22) collected in the Asian native area were also, and unexpectedly, found to be contaminated with *D. subpulchrella* individuals, namely CN-Nin with 7 *D. subpulchrella* individuals and to a lesser extent JP-Tok with 1 *D. subpulchrella* individual (both out of 50 individuals in total). More surprisingly, but confirming a gene-based analysis by D. Obbard (pers. comm.), the DE-Jen pool collected in Jena (Germany) was found to be contaminated with 5 to 6 *D. immigrans* individuals (out of 100). These observations may indicate that great care should be taken when analyzing sequencing data from wild-caught samples, and that more attention should probably be paid to species identification prior to sequencing. High-throughput metabarcoding and non-destructive approaches, such as those recently proposed by Piper *et al*. (28), may represent valuable alternatives to sometimes difficult morphological identification by allowing rapid and efficient diagnosis of *D. suzukii* samples at any life stage. Such efforts may be even more critical for Pool-Seq experiments, since filtering out contaminated sequences (e.g., using competitive mapping) is far more challenging than discarding mislabeled Ind-Seq samples, especially when the sample is contaminated by individuals from very closely related species (such as *D. subpulchrella* for *D. suzukii*).

Although two different *D. suzukii* genome assemblies were used to build the species-discriminating *k–mer* dictionary, all (pure) *D. suzukii* Ind-Seq and Pool-Seq samples showed a small but non-negligible fraction of their sequences (from 1.14% to 2.78%) assigned to *D. subpulchrella* by the most stringent criterion. Because i) the *D. suzukii* reference genome assemblies were derived from isofemale lines established from individuals sampled in the North American (5) and European (23) invaded areas; and ii) *D. subpulchrella* has not been yet described (to our knowledge) outside the Asian native range of *D. suzukii*; it is highly unlikely that this pattern is the result of pervasive gene flow between the two species, but rather can be explained by the close phylogenetic relationship between the two species. Indeed, some *D. subpulchrella*-discriminating *k–mers* may actually map to orthologous regions not represented in the *D. suzukii* reference assemblies and/or capture shared genetic variation between the two species due to incomplete lineage sorting (ILS). Including more reference assemblies (e.g., from different strains) for each target species may be considered as a valuable strategy to improve both the sensitivity (by ‘positive filtering’ of the discriminating *k–mers* that capture intraspecific genetic variation) and specificity (by ‘negative filtering’ of the incompletely sorted *k–mers*). The optimal number of representative assemblies is thus likely to both depend on the relatedness of the selected target species and for each target species on their genetic diversity. Alternatively, the misassigned short read sequences found in the analyzed samples can be included in the construction of the *k–mer* dictionary, assuming that the considered samples are not contaminated and are ‘pure’ representatives of the corresponding target species. Such refined target dictionaries may even further allow providing (rough) estimates of the genome-wide level of interspecific gene flow, or at least the identification of highly admixed individuals. Hence, in the sample of identified *D. subpulchrella* individuals, if about 2% of the short-read sequences were assigned to *D. suzukii* (in a similar but reversed pattern as observed for *D. suzukii* individuals), one (presumably) *D. subpulchrella* individual had nearly 10% of its sequences assigned to *D. suzukii*. The status of this sample may be of special interest for further study as it could represent a previously unreported case supporting some recent (i.e., only a few generations back) admixture events between *D. suzukii* and *D. subpulchrella*. As discussed by Lalyer *et al*. (17), if no such recent events have been reported to date, several studies suggest that hybridization has occurred between these two sister species (7).

Overall, the present analysis allowed the definition of a large curated dataset consisting of > 60 population samples representative of global genetic diversity, which may be valuable for further *D. suzukii* population genetics studies. Although constructed with the analysis of *D. suzukii* samples in mind, the *k–mer* dictionary developed here may be directly relevant to the analysis of the level of contamination of samples from other target species such as *D. simulans* or *D. melanogaster*. Likewise, the current dictionary also allows for the rapid identification of Wolbachia-infected samples, which may be of interest for a first rapid screening of drosophilids samples since the set of Wolbachia-discriminating *k–mers* was built by combining *D. simulans* and *D. melanogaster* Wolbachia assemblies. More generally, while we advocate careful sample identification and verification prior to sequencing, the proposed framework is straightforward and computationally efficient. It thus could be considered as a routine post-hoc quality check approach to be applied prior to any data analysis and prior to data submission to public repositories.

## Data availability

The CLARK and Clark-l*k–mer* databases and the (cleaned) assemblies used to build them have been made publicly available from the Data INRAE repository (12). The compressed archive also contains scripts used to run CLARK and Clark-lanalyses and parse the results. All sequencing data analyzed in this study are publicly available under the accession IDs reported in Tables 1, S2 and S3.

## Supplementary Materials

Supplementary Figures S1 to S5 are provided in the accompanying PDF file `Figures_S1_to_S5.pdf`. Supplementary Tables S1 to S5 are provided as Excel spreadsheets in the accompanying Excel file `Tables_S1_to_S5.xls`.

## Acknowledgements

I am grateful to the genotoul bioinformatics platform Toulouse Occitanie (Bioinfo Genotoul, https://doi.org/10.15454/1.5572369328961167E12) for providing computing resources. I also wish to thank Darren Obbard, Alan Bergland, Joaquin Nunez and Nicolas Rode for helpful discussion and feedback.

## Notes

https://doi.org/10.57745/HYTIBH

### Competing Interest Statement

The authors have declared no competing interest.

### Summary of Updates

This version of the manuscript has been revised to include the PCI Genomics badge following recommendation of the previous version (https://doi.org/10.24072/pci.genomics.100244). Line numbers have also been removed.

