## Supplementary Figures S1, S2, S3, S4 and S5 for "Efficient k-mer based curation of raw sequence data: application in *Drosophila suzukii*"

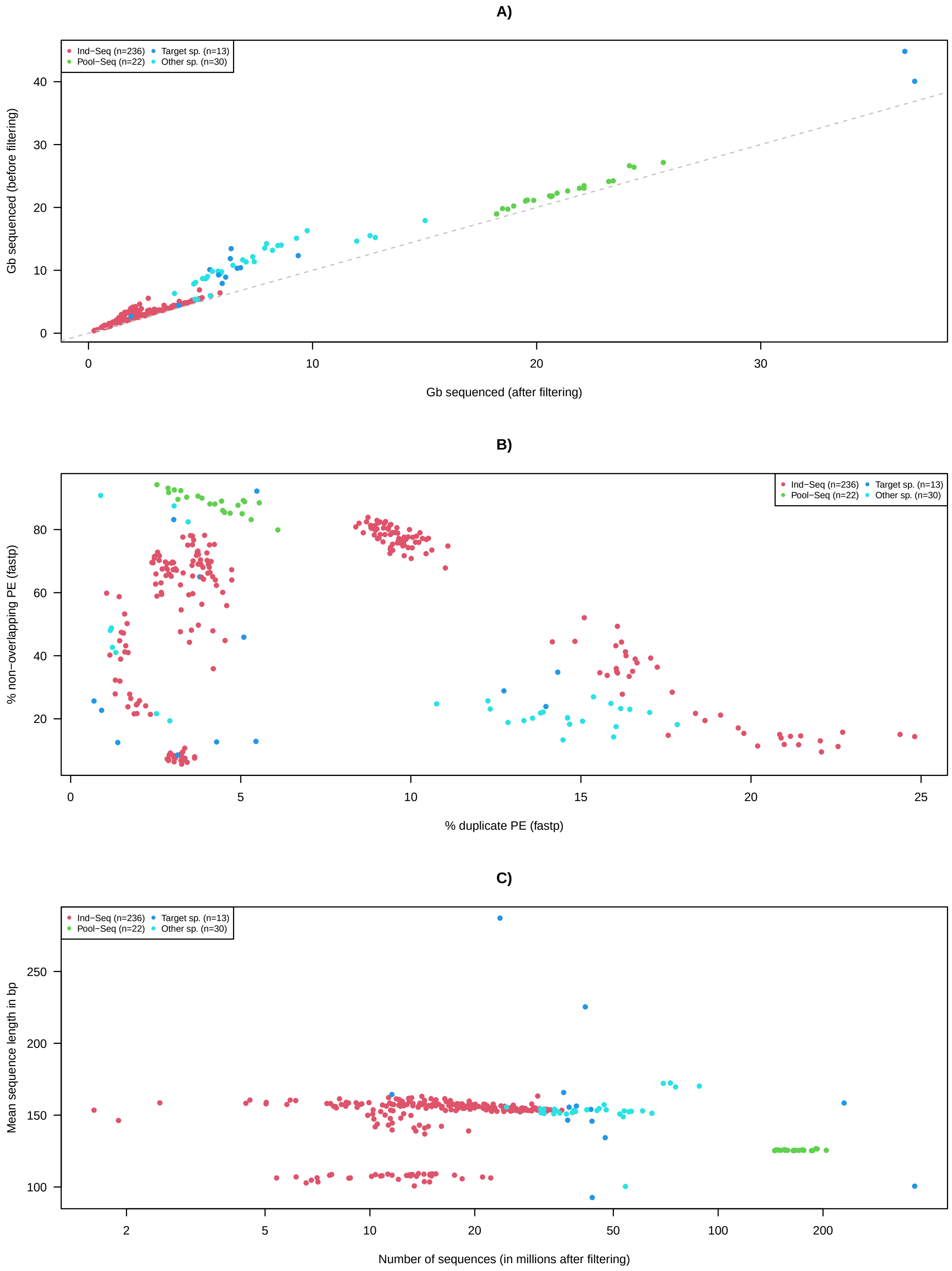


**Fig. S1.** Description of the query short-sequencing data consisting of 305 samples from i) 12 target species ('target sp.' in blue); ii) 31 additional drosophilid species ('other sp.' in light blue); iii) 236 *D. suzukii* individuals from Lewald *et al.* (18) ('Ind-Seq' in red); and iv) 22 *D. suzukii* pools of individuals from Olazcuaga *et al.* (22) ('Pool-Seq' in green). A) Number of sequenced bases before and after filtering with `fastp` (4). B) Estimated percentage (`fastp`) of duplicate (x–axis) and non-overlapping (y–axis) read pairs. C) Number and average sequence length in the different filtered data sets used for assignment analysis. See Tables S1 and S2 for more details.

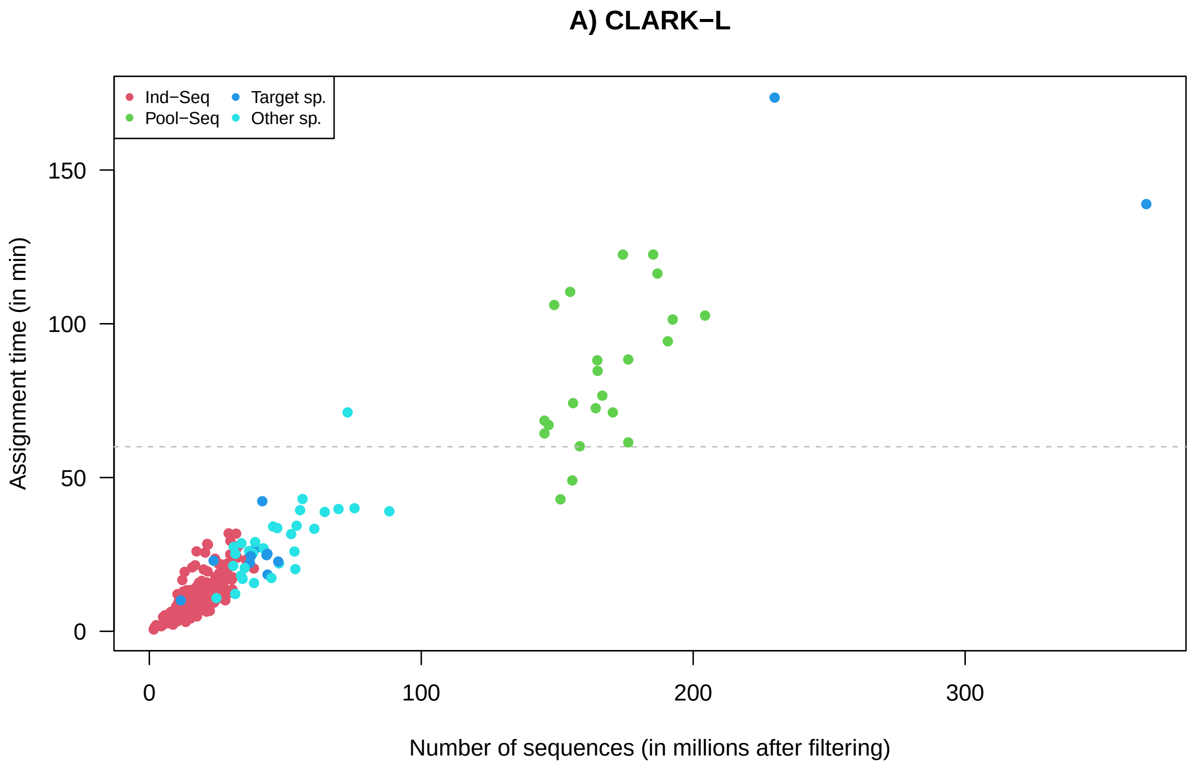


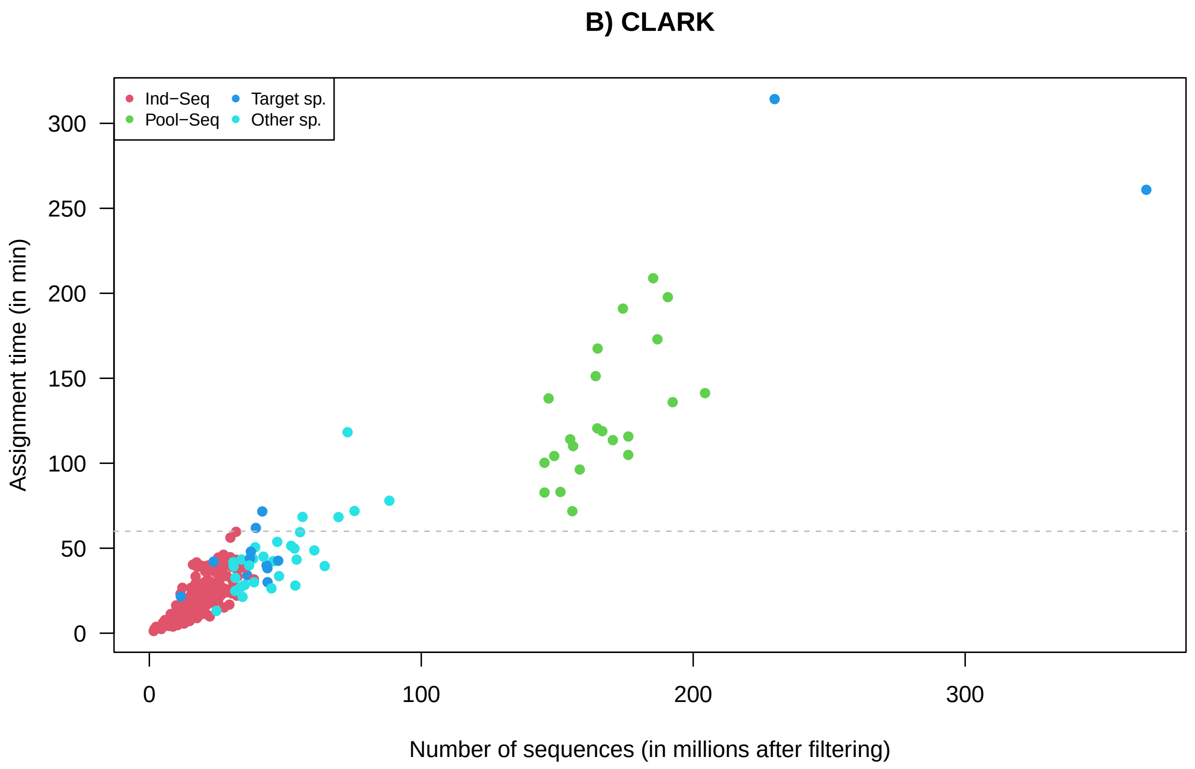


**Fig. S2.** Time spent for assigning all the sequences (i.e., excluding time for loading the k-mer dictionary) with CLARK-L (A) and CLARK (B) as a function of the number of sequences contained in the 301 whole-genome sequencing datasets. The gray dashed horizontal line represents 1 hour. Sequence length was representative of typical short-read datasets, with sample means ranging from 92.7 bp to 287 bp and a median size of 155 bp (Figure S1C).

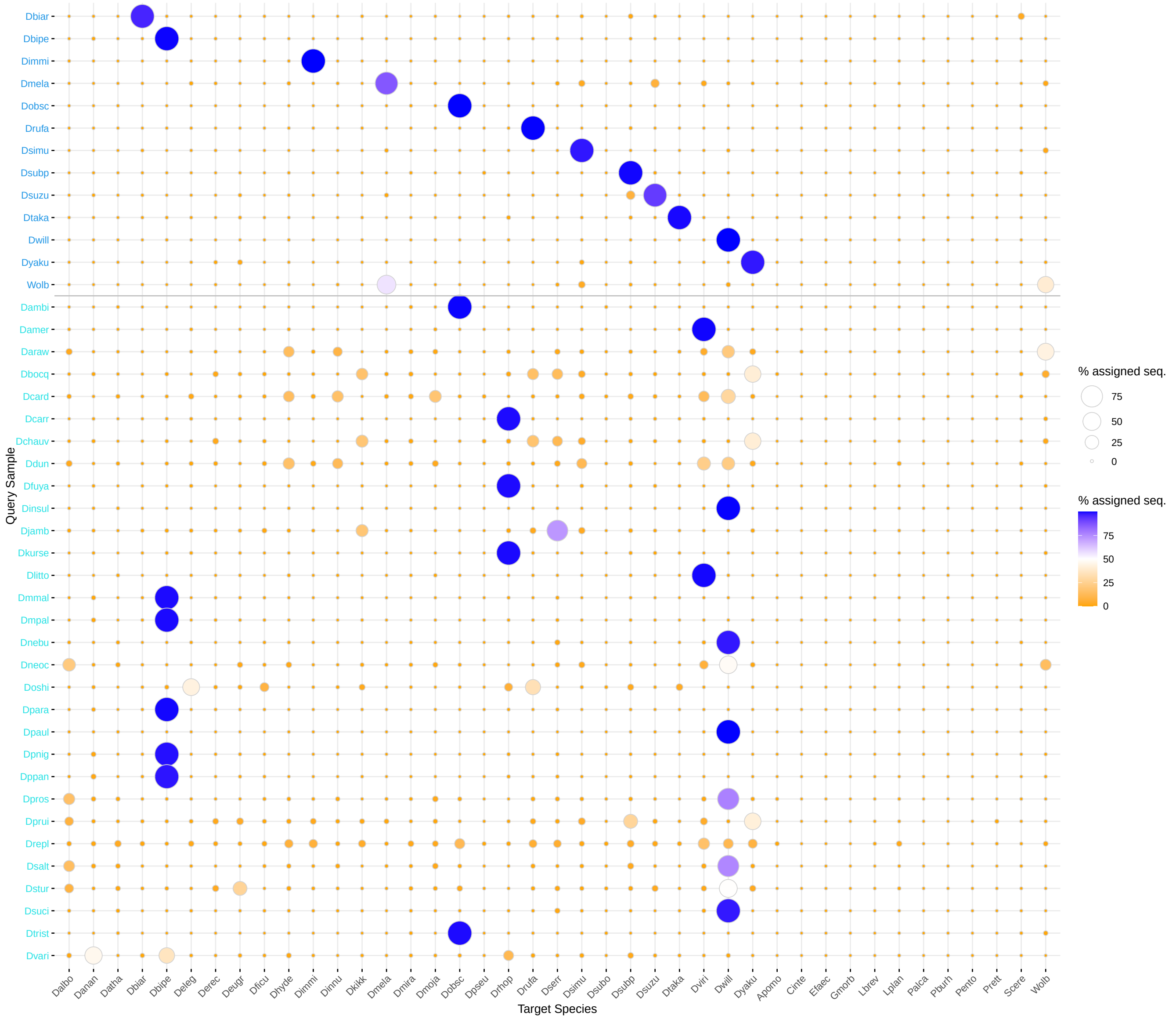


**Fig. S3.** Bubble plots summarizing assignment results obtained with CLARK-L using the most stringent sequence assignment criterion (i.e., $nk \geq 5$ and $c > 0.95$, see the main text) for 13 samples (labeled in dark blue at the top of the y-axis) belonging to species represented in the target *k–mer* dictionary and 30 other unrepresented drosophilid species. The 41 target species (29 drosophilid and 13 commensals or pathogens) are listed on the x-axis.

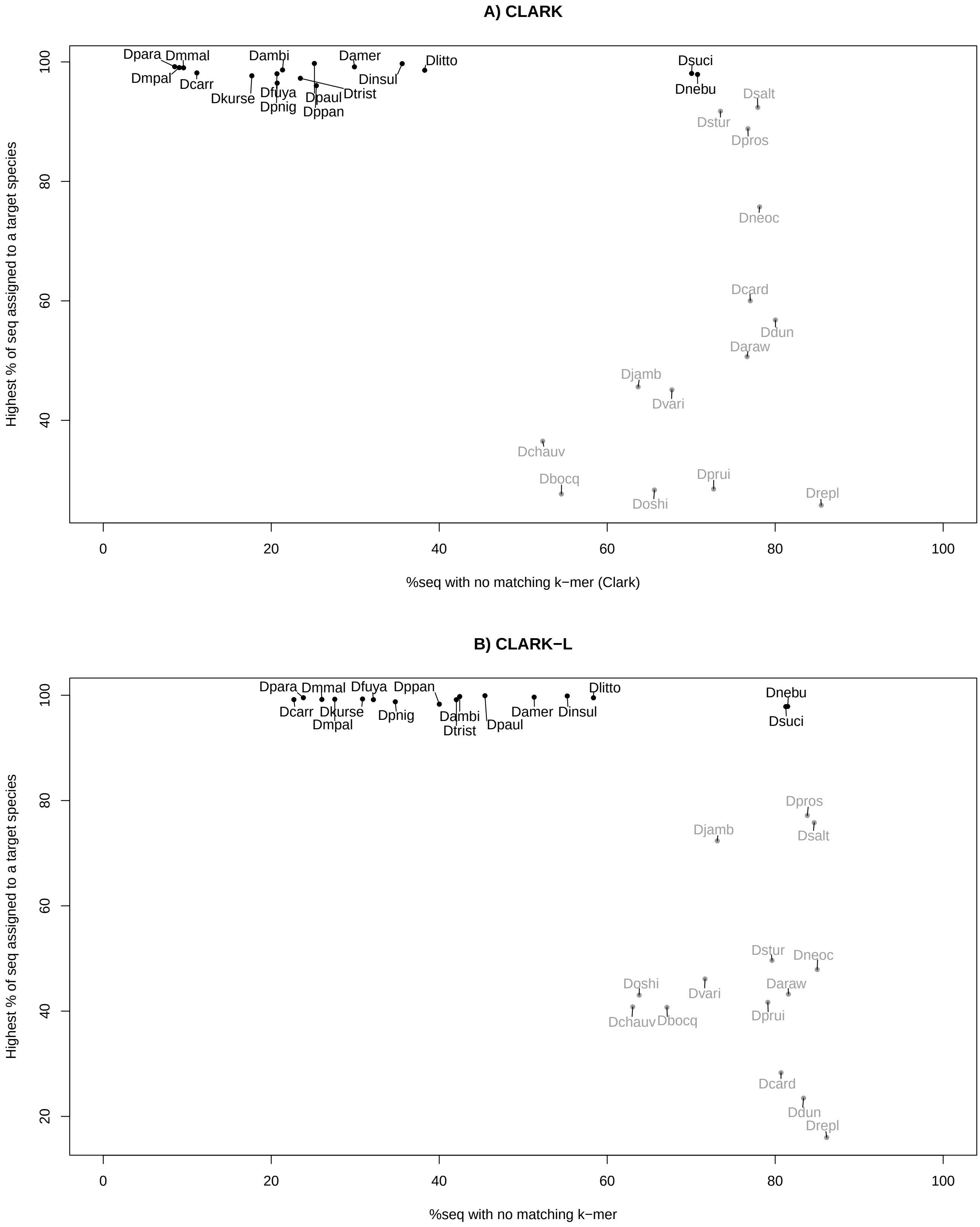


**Fig. S4.** Highest observed percentage of sequences assigned to a target species with CLARK (A) and CLARK-L (B) as a function of the percentages of sequences with no matching *k–mer* for the 30 samples belonging to non-target species (see Table S3 for details). Sequence assignment was performed using the most stringent criterion (i.e., $nk \geq 5$ and $c > 0.95$, see the main text). The 16 samples with >96% of their reads assigned to a given target species are represented in black, the 14 others are represented in grey.

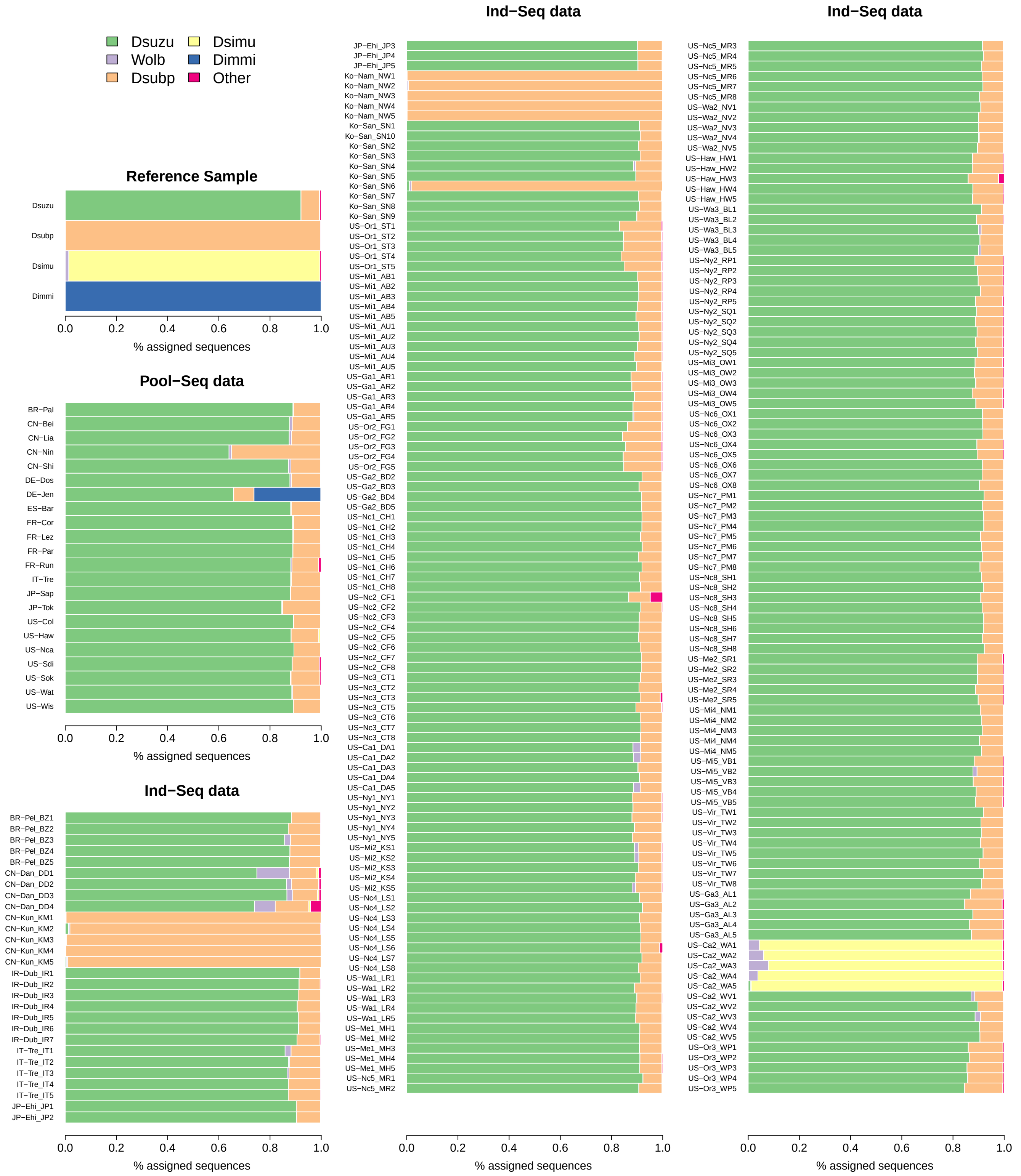


**Fig. S5.** Barplots summarizing assignment results obtained with CLARK-L using the most stringent sequence assignment criterion (i.e., $nk \geq 5$ and $c > 0.95$, see the main text) for the *D. suzukii* Ind-Seq (n=236) and Pool-Seq (n=22) samples. For each sample, the proportions of sequences assigned to the 5 target species that contribute at least 5% of the sequences of any of the 258 samples are shown using the color code indicated in the top left legend. The proportions of sequences assigned to the 36 other target species are shown in gray.
